# Structural, biophysical, and virological mechanistic characterization of HIV-1 capsid-targeting antivirals

**DOI:** 10.64898/2025.12.30.696960

**Authors:** Karen A. Kirby, William M. McFadden, Lei Wang, Haijuan Du, Huanchun Zhang, Andres Emanuelli Castaner, Zachary C. Lorson, Arvin Nafisi, Anastasia Selyutina, Charlotte Luchsinger, Atsuko Hachiya, Maria E. Cilento, Alexa A. Snyder, Shreya M. Ravichandran, Xinyong Cai, Philip R. Tedbury, Ashwanth C. Francis, Gregory B. Melikyan, Felipe Diaz-Griffero, Zhengqiang Wang, Stefan G. Sarafianos

**Author notes:** Corresponding Author: Stefan G. Sarafianos. State Key Laboratory of Fine Chemicals, Department of Pharmacy, School of Chemical Engineering, Dalian University of Technology, Dalian 116024, China (Lei Wang).

## Abstract

Due to its significant role in virus replication, the HIV capsid is an attractive antiviral target. This is validated by the recent clinical approval of lenacapavir for both treatment and pre-exposure prophylaxis (PrEP). PF74 is a well-characterized capsid-targeting antiviral that was discontinued in further study due to potency and metabolic issues. We hypothesized that making chemical modifications at certain sites of PF74 could result in capsid-targeting antivirals with improved potency and bioavailability. Our cumulative studies show that making changes at the R1 and R3 positions of PF74 results in compounds with increased antiviral potency, increased stability of wild-type HIV capsid hexamers and virions, tighter binding to wild-type HIV capsid hexamer compared to PF74, and different interactions at the “FG” binding site of capsid compared to PF74. These data provide insights into the design of future capsid-targeting antivirals relevant for clinical use.

## Introduction

The HIV-1 capsid plays a critical role in many steps of virus replication (1). At early stages of replication, after viral fusion with the host cell membrane, the capsid core is released into the cell and masks the viral nucleic acid from the host immune system defense response as it travels through the cytoplasm to the nucleus. The capsid core binds to several host factors, including CPSF6 and nucleoporins such as Nup358 and Nup153 that facilitate entry into the nucleus where uncoating of the core and reverse transcription of the viral RNA genome begins (2–6). The capsid core also directs the HIV-1 pre-integration complex to nuclear sites where the newly transcribed HIV-1 DNA is integrated into the host genome (7–10). At late stages of replication, after translation of the new viral proteins, the capsid domains of the Gag polyproteins interact with each other to help form immature viral particles at the membrane of the host cell. Once the immature particles are released from the cell, proteolysis of gag occurs, leading virion maturation, where the viral RNA genome is packaged into the assembled capsid core.

The HIV-1 capsid protein (CA) comprises an N-terminal domain (NTD) and a C-terminal domain (CTD). The CA protein assembles to form hexamers and pentamers, of which ∼250 and 12, respectively, further combine to form the fullerene-like mature capsid core. The mature capsid core has multiple binding sites that are important for interactions with cellular host factors that facilitate virus replication. The primary binding site in the HIV-1 capsid core is termed the “FG” binding pocket, since several host factors have a phenylalanine-glycine (“FG”) motif that binds at this pocket (11). This pocket is located between the NTD of one CA monomer and the CTD of a neighboring CA monomer within a hexamer (12). Given the biological importance of this pocket for entry into the host cell nucleus, it is a desirable antiviral drug target (13).

Over 30 million individuals are estimated to receive antiretroviral therapy (ART) (14), a well-tolerated co-formulation of multiple HIV-1 inhibitors that target various stages in the viral life cycle, decrease viral replication and transmission, and prevent the onset of acquired immunodeficiency syndrome (AIDS) (15–18). The long-acting and highly-potent drug lenacapavir (LEN, GS-6207, Sunlenca®), which binds at the “FG” binding pocket, is a first-in-class capsid-targeting antiviral approved for use in highly treatment-experienced (HTE) individuals as of 2022 and is currently the only capsid-targeting antiretroviral approved for clinical use (19–24). This validates the HIV capsid as a viable drug target. Other capsid-targeting antivirals, including PF74 that also targets the “FG” binding pocket, have also been well-studied for their effect on viral replication, but have been discontinued in clinical studies due to poor metabolic stability and potency (13, 25–30).

Our group and others have previously reported compounds derived from PF74 that overcome known issues associated with PF74 (13). We synthesized more than 300 PF74 analogs with chemical modifications at the R1 and R3 sites of PF74, and identified several compounds that increased the stability of WT CA hexamers in thermal shift assays, had better antiviral potency in cell-based assays, and improved metabolic stability compared to PF74 (31, 32). We hypothesized that chemical modifications in these compounds may lead to enhanced antiviral activity through significant changes in the efficiency of binding, as well as more pronounced structural effects. Here, we focus on additional biophysical and structural mechanistic studies on 10 of the most promising lead compounds (Figure 1), the most potent of which is ZW-1261 (EC_50_ = 0.022 µM). Together, these data enable structure-guided activity analysis to optimize future drug design at the clinically-targeted “FG” binding site.

**Figure 1.**
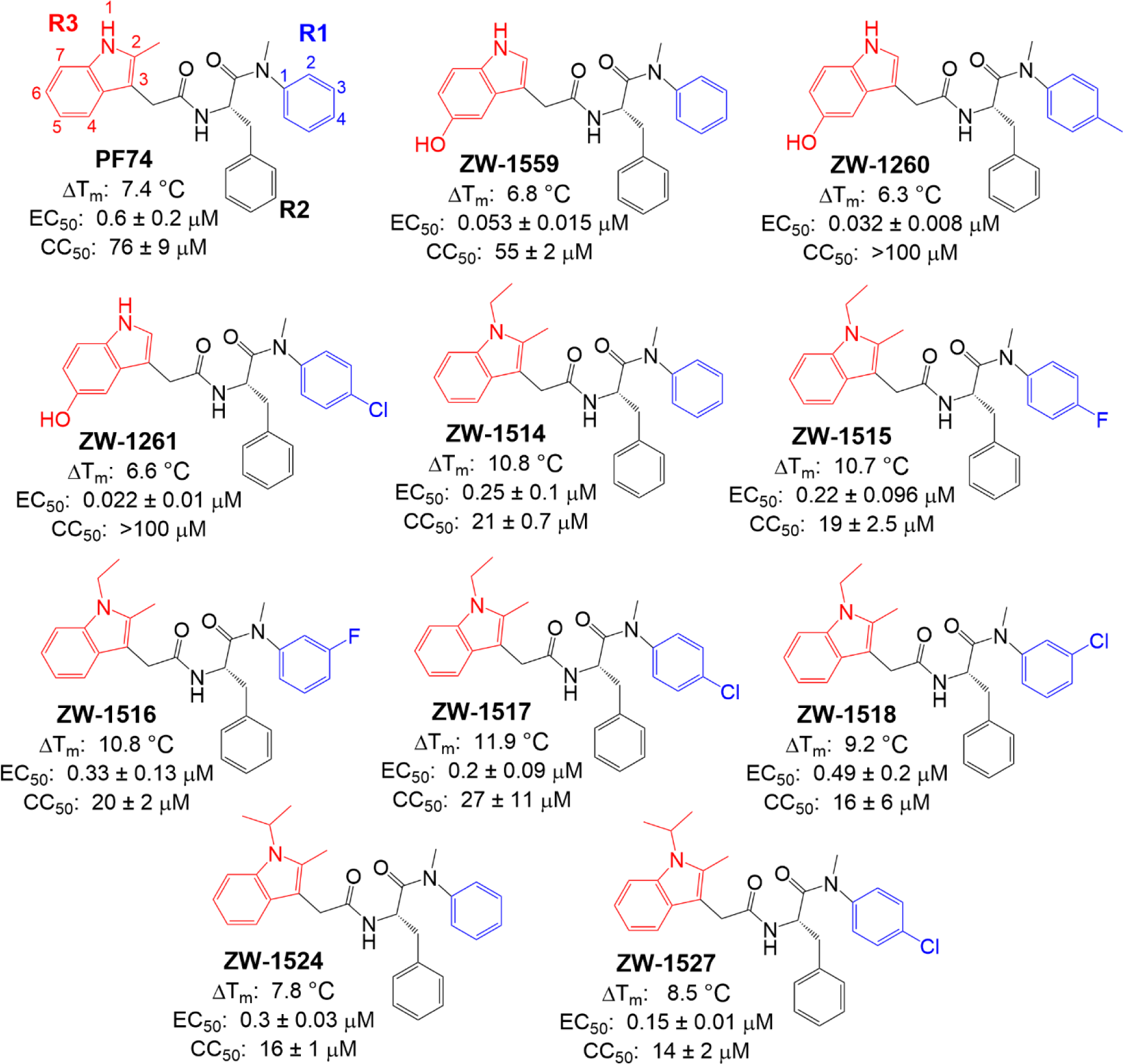
Chemical structures of PF74 and lead compounds that were determined to affect the stability of crosslinked CA hexamers and demonstrate antiviral activity against HIV-1. R1 regions are shown in blue; R3 regions are shown in red. R1 phenyl ring numbers and R3 indole ring numbers are shown on PF74. We previously reported their synthesis, thermal shift (ΔT_m_; using WT CA hexamers), antiviral (EC_50_; using WT HIV-1 virus and TZM-GFP cells), and cytotoxicity (CC_50_; using TZM-GFP cells) data (31, 32).

## Results

### Antiviral potency of PF74 analogs against different HIV-1 subtypes and HIV-2

We examined the antiviral activity of our two most potent lead compounds, ZW-1260 and ZW-1261 against different HIV-1 subtypes and HIV-2. Both ZW-1260 and ZW-1261 inhibited HIV-1 clades B, C, AE, and AG, with ZW-1261 being more potent in most cases than ZW-1260 (EC_50_s for ZW-1261 ranged from 0.018 to 0.060 µM against HIV-1 subtypes, Table 1). Both compounds were more potent than PF74, with ZW-1261 being ∼16-fold more potent against subtype B, ∼10-fold more potent against subtype C, ∼28-fold more potent against subtype AE, and ∼14-fold more potent against subtype AG (Table 1). Both compounds were also more potent than PF74 against HIV-2, with ZW-1261 being ∼7-fold more potent than PF74 (Table 1). These data demonstrate that lead compounds ZW-1260 and ZW-1261 can efficiently inhibit non-B HIV-1 subtypes.

**Table 1.**
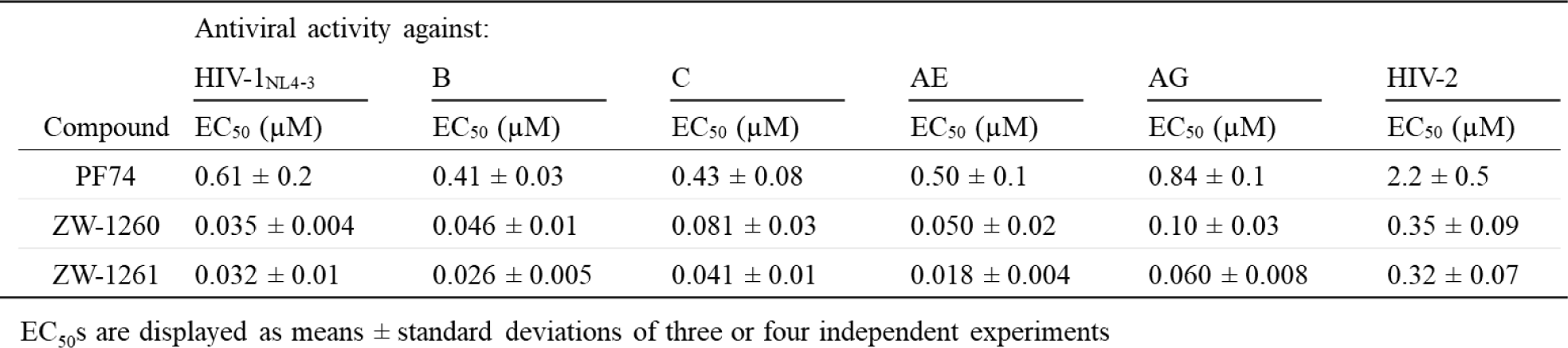
Cell-based antiviral activity assays of PF74, ZW-1260, and ZW-1261 against HIV-1 subtype B, C, AE, and AG, and HIV-2.

### Effects of compounds on early vs. late stages of HIV-1 replication

We tested the 10 lead compounds (Figure 1) for their effect on early *vs.* late stages of HIV-1 replication. Early-stage events (virus entry through integration) are examined by using pseudotyped HIV-1 virus with a vesicular stomatitis virus glycoprotein (VSV-G) envelope to generate a single round of HIV-1 infection (25). Late-stage events (steps following integration, including assembly and maturation) are studied by packaging virus in the presence of compounds, and testing infectivity of the resulting viral supernatants in TZM-GFP target cells in the absence of the compounds (25). As expected, the control integrase strand transfer inhibitor (INSTI) raltegravir (RAL) inhibited early-stage, but not late stage, replication (Table 2). The control protease inhibitor nelfinavir (NFV) inhibited late-stage, but not early-stage, replication events (Table 2). Parent compound PF74 inhibited both early and late stages of viral replication, as previously reported (25). All 10 lead compounds also inhibited both early and late stages of replication similar to PF74 (Table 2).

**Table 2.** Antiviral activity of compounds in early and late stages of infection.

| Compound | Antiviral activity against: |  | Stage of action |
| --- | --- | --- | --- |
|  | VSV-G Pseudovirus | Production/Infection |  |
|  | EC<sub>50</sub> (μM) | EC<sub>50</sub> (μM) |  |
| PF74 | 0.52 ± 0.07 | 2.1 ± 0.6 | Early/Late |
| ZW-1260 | 0.033 ± 0.005 | 0.38 ± 0.09 | Early/Late |
| ZW-1261 | 0.021 ± 0.008 | 0.17 ± 0.02 | Early/Late |
| ZW-1514 | 0.26 ± 0.04 | 0.45 ± 0.06 | Early/Late |
| ZW-1515 | 0.22 ± 0.006 | 0.54 ± 0.03 | Early/Late |
| ZW-1516 | 0.43 ± 0.03 | 0.46 ± 0.2 | Early/Late |
| ZW-1517 | 0.16 ± 0.05 | 0.67 ± 0.2 | Early/Late |
| ZW-1518 | 0.47 ± 0.01 | 0.78 ± 0.1 | Early/Late |
| ZW-1524 | 0.39 ± 0.04 | 1.2 ± 0.2 | Early/Late |
| ZW-1527 | 0.33 ± 0.03 | 0.74 ± 0.07 | Early/Late |
| ZW-1559 | 0.057 ± 0.005 | 0.38 ± 0.07 | Early/Late |
| RAL | 0.02 | >1.5 | Early |
| NFV | >20 | 0.02 | Late |

<sup>a</sup>EC<sub>50</sub>s are displayed as means ± standard deviations of three separate experiments.

### Effect of select lead compounds on HIV-1 capsid core stability

We next assessed how lead compounds influence the stability of native HIV-1 capsids. To do this, we employed a well-established capsid stability assay that monitors the loss of a high-avidity capsid-binding chimeric Cyclophilin A–DsRed (CypA-DsRed, CDR) fusion protein using time-resolved confocal imaging (26). The CDR marker efficiently labels HIV-1 cores, and its fluorescence loss correlates with the capsid lattice disassembly in permeabilized virions (33–35). In the DMSO vehicle control, CDR signal decreased over time, indicating progressive loss of capsid lattice stability (Figure 2). In contrast, treatment with 10 µM lead compounds ZW-1261, ZW-1559, ZW-1514, ZW-1517, and ZW-1524 maintained nearly 100% CDR signal for up to 1 hour, demonstrating the stabilization effect of these lead compounds on native HIV-1 capsids.

**Figure 2.**
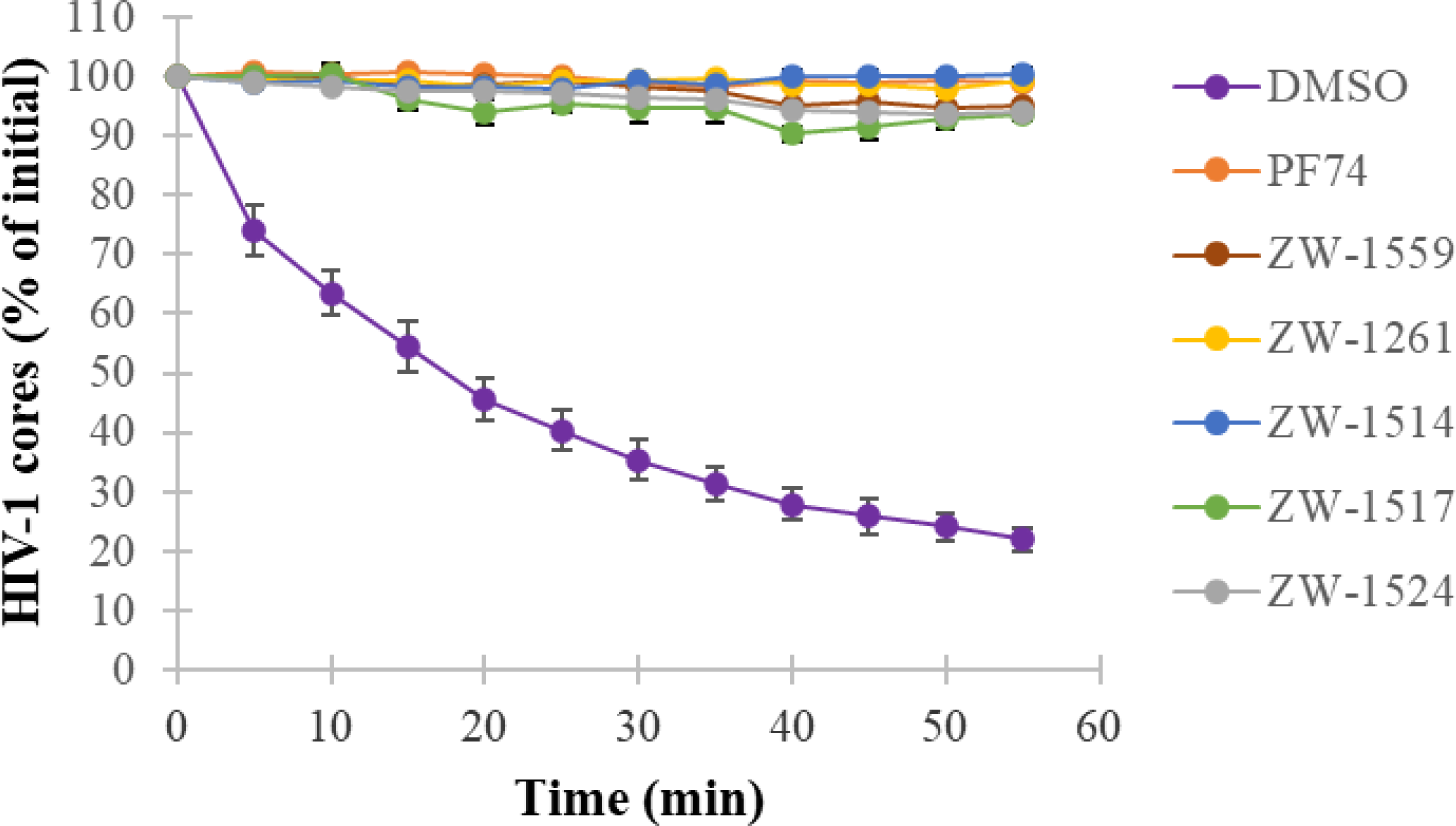
Kinetics of single-virus core-stability, as measured by the loss of CypA-DsRed bound to viral membrane permeabilized HIV-1 cores *in vitro*. The DMSO vehicle control and 10 μM of indicated compound was added to CypA-DsRed bound cores and the loss of the fraction of CypA-DsRed cores was quantified by confocal imaging. The HIV-1 cores were stabilized in the presence of PF74 and its analogs compared to the respective half-time of the intrinsic core-stability (∼13 mins) observed in control DMSO samples. Data are averaged from four independent experiments.

We additionally tested the effect of lead compound ZW-1261 on CA stability using a fate-of-the-capsid assay (36–40). This fluorescent western blot assay can be applied at any time during HIV-1 infection to determine the relative amount of capsid monomers and small lattice fragments (which are soluble) compared to the amount of cores (which are pelletable). The input sample is shown in Figure 3A. PF74 was used as a control, and at 10 µM results in primarily soluble CA (Figure 3B), indicating destabilization of the core in this assay. At low concentrations, ZW-1261 destabilizes the core (0.5-1 µM) like PF74; however, at higher concentrations (2.5-5 µM), stabilization of the core was observed (Figure 3C), suggesting a possible dual-phase inhibition mechanism.

**Figure 3.**
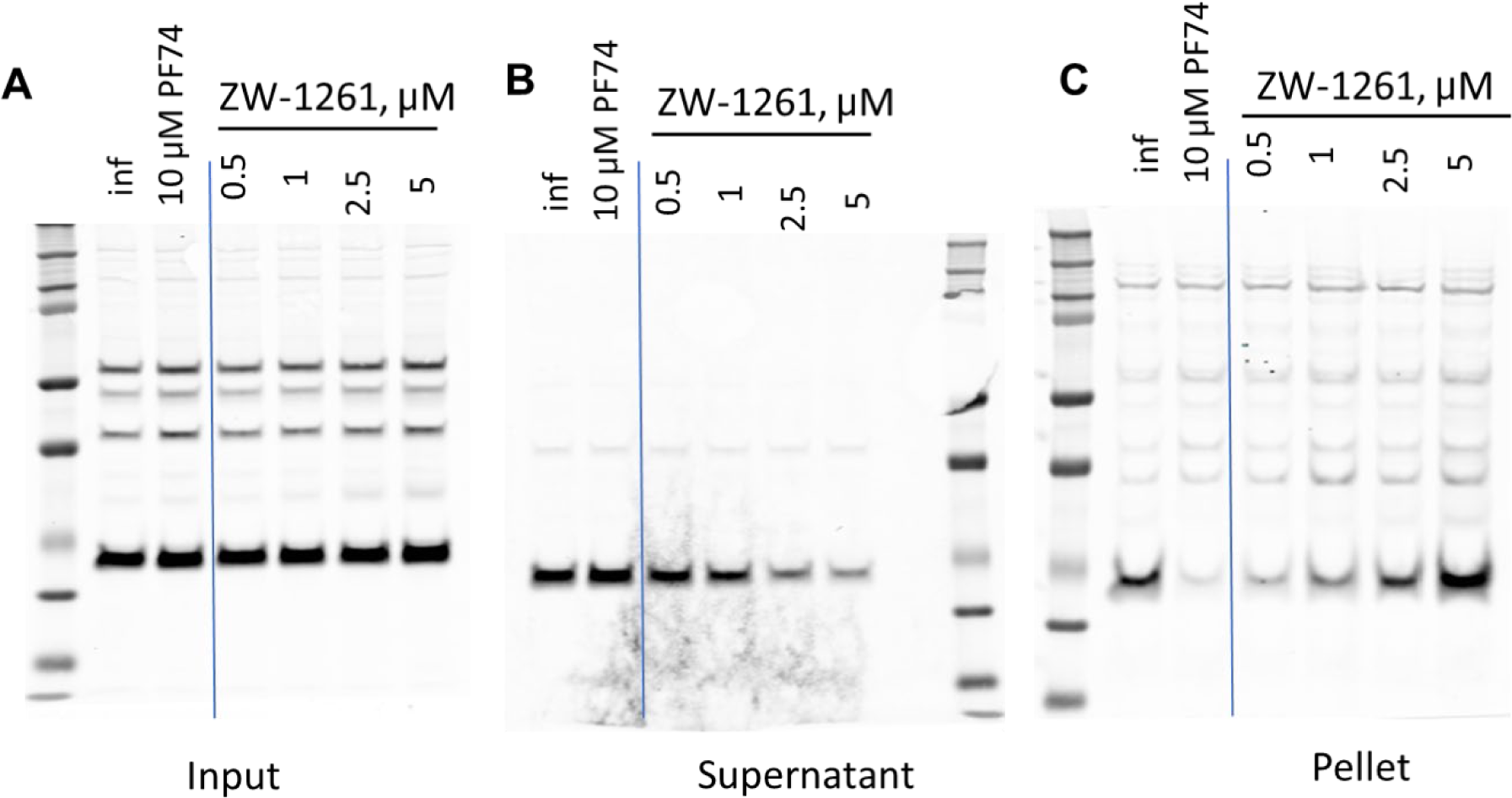
Fate-of-the-capsid assay with ZW-1261. (**A**) Input sample prior to centrifugation. (**B**) Supernatant (unassembled CA) and (**C**) pellet (assembled CA, including cores) after centrifugation. 10 µM PF74 was used as a control; at this concentration PF74 results primarily in soluble, unassembled CA. The lowest band in (**B**) decreases with increasing concentration of ZW-1261, indicating that soluble, unassembled CA is decreasing. The lowest band in (**C**) is increasing with increasing concentration of ZW-1261, indicating stabilization of assembled CA.

### Effect of lead compound ZW-1261 on HIV-1 nuclear import

We also tested the effect of lead compound ZW-1261 on the nuclear import of HIV-1 capsid using a fluorescence-based western blot assay as previously described (40, 41). The PF74 control prevents the nuclear import of capsid (Figure 4A *vs.* 4B). We confirmed the location and purity of cellular fractions using anti-tubulin and anti-Nup153 antibodies, which are cytosolic and nuclear markers, respectively (Figures 4C,D). At low concentrations of ZW-1261 there is a small decrease in the entry of capsid into the nucleus. However, at high concentrations of ZW-1261, there is an increase in the entry of capsid into the nucleus (Figure 4A *vs.* 4B). This correlates an increase of core stability with entry of capsid into the nucleus.

**Figure 4.**
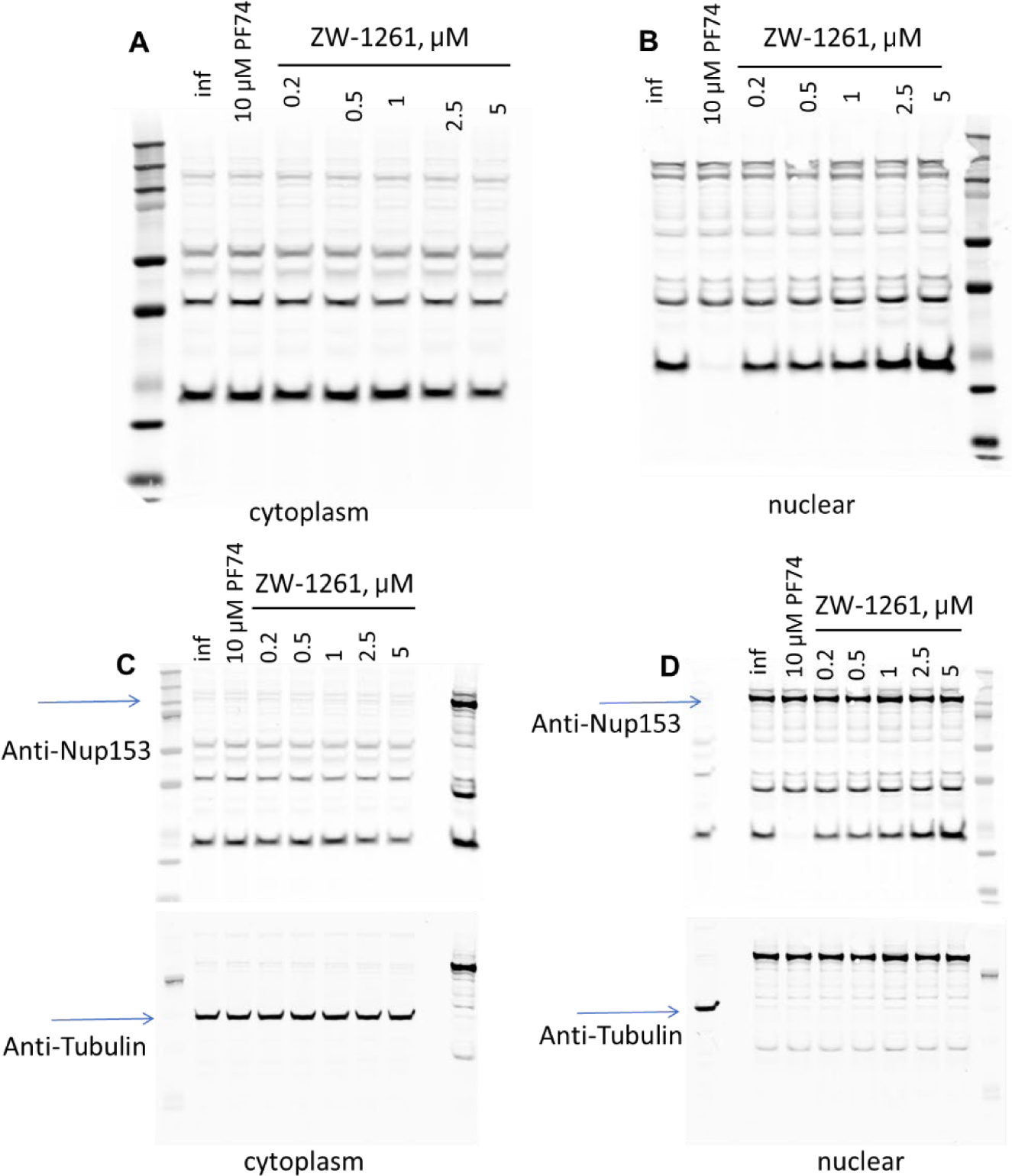
Effect of ZW-1261 on nuclear import. Cells infected with HIV-1 in the presence of PF74 control or lead compounds are separated into cytosolic (**A**) and nuclear (**B**) fractions. These fractions were analyzed by western blotting using anti-p24 antibodies. To verify the location and purity of cellular fractions, anti-tubulin and anti-Nup153 antibodies, which are cytosolic (**C**) and nuclear markers (**D**), respectively, were also used.

### Effect of lead compound ZW-1261 on CPSF6 aggregation in nuclear speckles

To further investigate the mechanism of action of ZW-1261 during replication, we examined its effect on the formation of CPSF6 aggregates that colocalize with nuclear speckles during HIV-1 infection (40, 42, 43). We found that when ZW-1261 is present during infection, it blocks the HIV-1–induced translocation of CPSF6 to nuclear speckles, similar to PF74 and lenacapavir precursor GS-CA1 (Figure 5A) (43). Notably, when ZW-1261 is added after infection, once CPSF6 has already re-localized to nuclear speckles, it actively disassembles CPSF6 from these compartments, mimicking the effect of PF74 but not GS-CA1 (Figure 5B).

**Figure 5.**
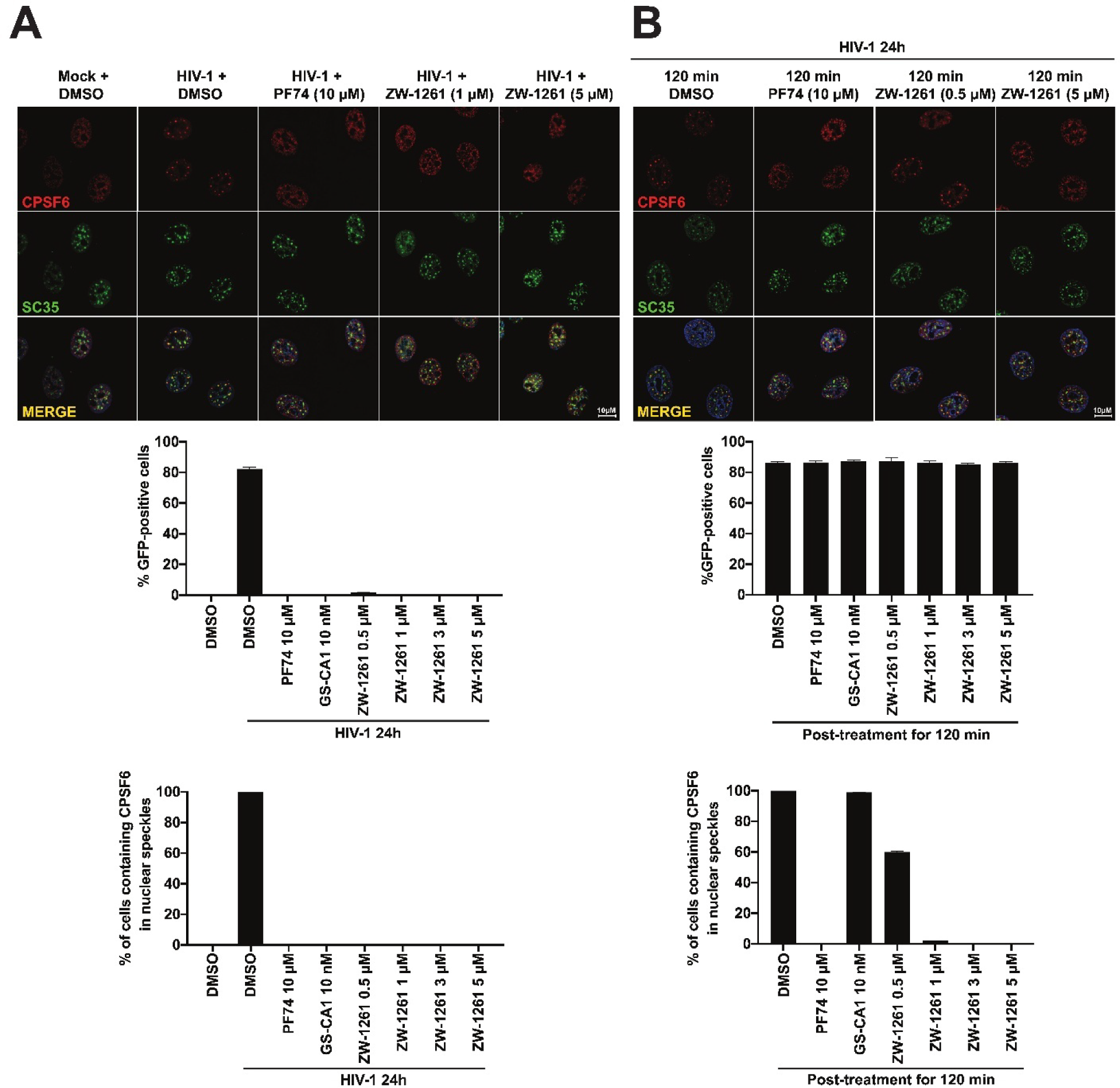
ZW-1261 disaggregates preformed CPSF6 complex in nuclear speckles by HIV-1 infection. (**A**) Human A549 cells were infected with HIV-1-GFP at an MOI of 2 for 24 h in the presence of 10 µM PF74, 10 nM GS-CA1, increasing concentrations of ZW-1261 (0.5 to 5 µM) or DMSO (vehicle control). Infection was assessed as the percentage of GFP-positive cells by flow cytometry as shown in the graphs. (**B**) To induce the translocation of CPSF6 to nuclear speckles, A549 cells were challenged with HIV-1-GFP at an MOI of 2 for 24 h. Subsequently, cells were incubated for 120 min with 10 µM PF74, 10 nM GS-CA1, increasing concentrations of ZW-1261 (0.5 to 5 µM) or DMSO (vehicle control). Infection was assessed as the percentage of GFP-positive cells by flow cytometry as shown in the graphs. For (**A**) and (**B**) cells were fixed, permeabilized, and coimmunostained using rabbit polyclonal antibody to CPSF6 (red) with mouse monoclonal antibody against SC35 (pseudo-green, marker for nuclear speckles). Secondary antibodies were Alexa-594-conjugated donkey anti-rabbit IgG; and Alexa-647-conjugated donkey anti-mouse IgG, respectively. Nuclei were counterstained with DAPI (blue). Merging the red, green and blue channels generated the third image in each column; yellow indicates overlapping localization of the red and green channels. Scale bar, 10 µm. Percentage of cells containing CPSF6 in nuclear speckles (SC-35-positive compartments) was quantified from three independent experiments by visual examination of 200 cells per condition.

### Effect of substitutions on binding efficiency to WT HIV CA hexamer

We used biolayer interferometry (BLI) to assess the binding kinetics of select lead compounds to crosslinked HIV CA hexamers containing a C-terminal His tag. We immobilized His-tagged WT HIV-1 CA hexamers onto HIS1K Octet biosensors and examined the binding and dissociation of select analogs at 5, 10, and 20 µM using a previously described protocol (44, 45). The LEN control bound tightly to the crosslinked WT HIV-1 CA hexamers with a K_D_ of 0.89 nM (Table 3). This is primarily due to its ability to remain tightly bound to the CA hexamers as determined by its exceptionally slow dissociation rate (*k_off_* = 4.4 × 10^-5^ s^-1^). The PF74 control bound to the crosslinked WT HIV-1 CA hexamers with a K_D_ of 365 nM. PF74 demonstrated a slightly longer association rate than LEN (4.9 × 10^4^ M^-1^s^-1^ *vs.* 9.7 × 10^4^ M^-1^s^-1^, respectively) but had a much faster dissociation rate (3.5 × 10^-2^ s^-1^). Compounds ZW-1559, ZW-1260, and ZW-1261 which all have a 5-hydroxy group and lack a 2-methyl on the R3 indole ring all have lower K_D_ values than PF74, with ZW-1261 having a slower dissociation rate than PF74, ZW-1559, and ZW-1260 (Table 3). Compounds ZW-1514 and ZW-1517 that have an N1-ethyl group also have lower K_D_ values than PF74, with ZW-1517 having a slower dissociation rate than PF74 and ZW-1514 (Table 3). Compound ZW-1524, which has an N1-isopropyl group, has a higher K_D_ value than PF74 and the other ZW analogs tested (Table 3).

**Table 3.** Binding of select capsid-targeting compounds to crosslinked WT HIV CA hexamers measured by BLI. Binding constants (K_D_), rates of association (*k_on_*) and rates of dissociation (*k_off_*) are provided for the binding of listed compounds to crosslinked WT CA hexamers with C-terminal His tags. Data are averages of duplicates in at least two independent experiments.

| **Compound binding to**<br>**HIV-1 CA Hexamer** | **$K_D$ (nM)** | **$k_{on}$ ($\text{nM}^{-1}\text{s}^{-1}$)** | **$k_{off}$ ($\text{s}^{-1}$)** |
| --- | --- | --- | --- |
| LEN | $0.89 \pm 0.02$ | $4.9 \times 10^{-5} \pm 0.05$ | $4.4 \times 10^{-5} \pm 5.9 \times 10^{-7}$ |
| PF74 | $365 \pm 16.7$ | $9.7 \times 10^{-5} \pm 0.4$ | $3.5 \times 10^{-2} \pm 3.3 \times 10^{-4}$ |
| ZW-1559 | $157 \pm 9.6$ | $7.6 \times 10^{-5} \pm 0.5$ | $1.2 \times 10^{-2} \pm 1.8 \times 10^{-4}$ |
| ZW-1260 | $128 \pm 4.9$ | $1.7 \times 10^{-4} \pm 0.6$ | $2.1 \times 10^{-2} \pm 1.2 \times 10^{-4}$ |
| ZW-1261 | $105 \pm 2.1$ | $6.0 \times 10^{-5} \pm 0.1$ | $6.4 \times 10^{-3} \pm 4.3 \times 10^{-5}$ |
| ZW-1514 | $160 \pm 4.5$ | $6.7 \times 10^{-5} \pm 0.2$ | $1.1 \times 10^{-2} \pm 7.4 \times 10^{-5}$ |
| ZW-1517 | $71.6 \pm 1.8$ | $2.9 \times 10^{-5} \pm 0.05$ | $2.1 \times 10^{-3} \pm 4.0 \times 10^{-5}$ |
| ZW-1524 | $599 \pm 15.5$ | $2.9 \times 10^{-5} \pm 0.07$ | $1.7 \times 10^{-2} \pm 1.2 \times 10^{-4}$ |

### Effect of R3 substitutions on binding at the “FG”-binding pocket

To understand the specific molecular interations of lead compounds with HIV-1 CA, we determined X-ray crystal structures of our wild-type (WT) full-length CA construct (CA FL) in complex with 3 lead compounds that differ primarily in their R3 substitutions: ZW-1261 at 2.7 Å resolution (2-H and 5-OH R3 substitutions; Figure 6A), ZW-1514 at 2.8 Å resolution (2-methyl and N1-ethyl R3 substitutions; Figure 6B), and ZW-1527 at 2.5 Å resolution (2-methyl and N1-isopropyl R3 substitutions; Figure 6C). All three compounds bind at the “FG”-binding pocket, where host factors, PF74, and LEN also bind. This pocket is at the interface between the N-terminal domain of one CA monomer and the C-terminal domain of a neighboring CA monomer within a hexamer (12, 28, 29). ZW-1261 is able to make H-bond interactions with its 5-OH group with the C-terminal domain of the neighboring CA through direct interaction with the side chain of K182 and through interaction of an organized water molecule that bridges to Q179 (Figure 6A). These are new interactions with the C-terminal domain of the neighboring CA monomer within a hexamer that the parent PF74 compound is not capable of doing. ZW-1514, like PF74, does not interact with the C-terminal domain of the neighboring CA monomer other than the cation-π interaction between the indole ring and R173. The N1-ethyl substitution of ZW-1514 prevents the H-bond between N1 and Q63 that exists in the PF74 structure. This creates a slight steric effect and causes the R3 position to shift lower than PF74, creating a hydrophobic interaction bewteen the N1-ethyl group and the main chain Cα of Q67 (Figure 6B), which is a new interaction compared to PF74. ZW-1527 also does not interact with the C-terminal domain of the neighboring CA monomer. The N1-isopropyl group of ZW-1527 creates such a steric clash with Q63 that the entire R3 indole ring surprisingly flips ∼90° compared to PF74. This results in the N1-isopropyl group pointing toward the C-terminal domain of the neighboring CA monomer (Figure 6C).

**Figure 6.**
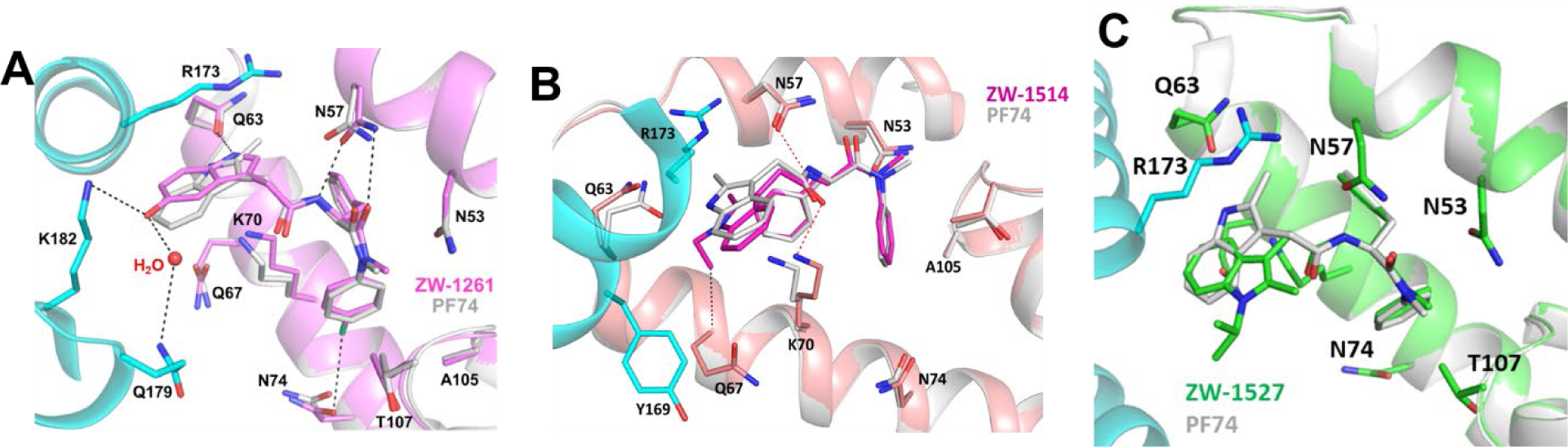
X-ray crystal structures of lead compounds with WT CA FL: ZW-1261 at 2.7 Å (**A**), ZW-1514 at 2.8 Å (**B**), and ZW-1527 at 2.5 Å (**C**). The WT CA FL N-terminal domain of one subunit is colored as dark pink cartoon for the ZW-1261 complex (**A**), light pink cartoon in (**B**), and green cartoon in (**C**). The C-terminal domain of the neighboring CA monomer is colored as cyan cartoon. ZW-1261 is shown as dark pink sticks in (**A**), ZW-1514 as light pink sticks in (**B**), and ZW-1527 as green sticks in (**C**). Residues around the “FG”-binding pocket are shown as colored sticks. Each new structure has been superposed to WT CA FL in complex with PF74 (PDB ID: 4XFZ), with the N-terminal domain shown as light gray cartoon and PF74 as light gray sticks.

## Discussion

LEN is a pioneering addition to the ART arsenal as a first-in-class capsid-targeting antiviral with sub-nanomolar potency that is formulated as a long-acting injectable therapy administered twice yearly for the maintenance of HIV suppression (13, 16, 19, 21–24, 46). LEN has also produced exciting preliminary results in pre-exposure prophylaxis (PrEP) trials, suggesting its promise for use in prevention of HIV-1 infection (24, 47, 48) and leading to its recent approval for PrEP as Yeztugo® (49, 50). However, the large size and complex chemistry of LEN complicate a structure-guided design approach (46, 51). By optimizing the PF74 scaffold, we were able to characterize the effects of chemical modifications on targeting the capsid to block HIV-1 replication.

In previous studies, we produced a range of PF74 derivatives by modification at the R1 and R3 regions (blue and red regions, respectively, Figure 1). Initial screening identified analogs that increased the stability of WT CA hexamers and were more potent than PF74 in cell-based antiviral assays (31, 32). Compounds ZW-1559, ZW-1260, and ZW-1261 differ from PF74 primarily in their R3 region, with the 2-methyl group removed and the addition of a 5-hydroxy group. These changes resulted in a significant (ranging from 10-to 27-fold) increase in potency against WT HIV-1 compared to PF74 (EC_50_ values of ZW-1559, ZW-1260, and ZW-1261 compared to PF74, Figure 1). Additional changes at the R1 region, including a 4-methyl or 4-chloro group further increased the antiviral potency against WT virus (EC_50_ value of ZW-1559 compared to ZW-1260 and ZW-1261, Figure 1). Compounds ZW-1514 through ZW-1518 differ from PF74 primarily by an ethyl group on the N1 position of the R3 region. This change results in an approximately 2-fold increase in antiviral potency compared to PF74 (EC_50_ of ZW-1514 compared to PF74, Figure 1). The inclusion of 4-fluoro or 4-chloro substitutions in the R1 region provides a slight increase in antiviral potency (EC_50_s of ZW-1515 and ZW-1517 compared to ZW-1514, Figure 1). 3-fluoro or 3-chloro substitutions decreased the antiviral potency (EC_50_s of ZW-1516 and ZW-1518 compared to ZW-1514, Figure 1). Compounds ZW-1524 and ZW-1527 differ from PF74 primarily by an isopropyl substitution on the N1 position of the R3 region. This change also results in increased antiviral potency compared to PF74 (EC_50_s of ZW-1524 vs. PF74, Figure 1). Like the other lead compounds, addition of a 4-chloro group in the R1 region increased antiviral potency (EC_50_ of ZW-1524 compared to ZW-1527, Figure 1). Overall, these results demonstrate that changes in the R3 region at positions 1 and 5, in combination with a 4-chloro group in the R1 region, can improve antiviral potency and increase stability of the HIV CA hexamer. In the current study, we examined these compounds to evaluate how different chemical modifications affect binding to CA and stability.

In addition to being potent inhibitors of subtype B HIV-1, compounds ZW-1260 and ZW-1261 can efficiently inhibit non-B HIV-1 subtypes (Table 1, Figure 1); we hypothesize that this is primarily due to the 5-hydroxy addition to the R3 region of the PF74 scaffold, with the 4-chloro group in the R1 region providing additional antiviral potency against most non-B HIV-1 subtypes. There is high sequence conservation in the HIV-1 CA, particularly in the “FG”-binding pocket (52). Antiviral activity against HIV-2 for PF74, ZW-1260, and ZW-1261 is ∼4-10-fold-worse than against HIV-1 (Table 1). Notably, LEN is also 8-10-fold less potent against HIV-2 than HIV-1 (19, 21). This is likely because of differences in sequence between HIV-1 and HIV-2 capsid at the “FG”-binding pocket (53). Specifically, at residue 70, which is in the middle of the “FG”-binding pocket and has key interactions with LEN, PF74, and related analogs. In HIV-1 residue 70 is a K, whereas in HIV-2, residue 70 is an R which has a longer side chain that could hinder access to the “FG”-binding pocket (54). Another difference in sequence between HIV-1 and HIV-2 capsid is at residue 179, which is part of the CTD and also involved in LEN binding. In HIV-1, residue 179 is a Q, whereas in HIV-2, the same residue is a P, which would reduce interactions with LEN compared to HIV-1 (54).

The mechanism of inhibition by LEN is under active investigation as it is complex, impacting both the early and late stages of HIV-1 replication (13, 19, 35, 55, 56). In the mature capsid, LEN disrupts the core assembly by binding to the functionally important “FG” binding pocket that natively interacts with several host proteins, like Nup153 or CPSF6 (11, 19, 29, 34, 35, 56–58). The mechanism of PF74 inhibition has been extensively studied and the compound has been shown to block both early and late stages in HIV-1 replication cycle (25). During early stages of HIV-1 infection, PF74 suppresses viral replication through a concentration-dependent bimodal mechanism of action (59). High concentrations of PF74 (10 μM) trigger early uncoating, affecting reverse transcription and reducing the amount of newly-synthesized viral DNA (25–30). Low concentrations (<2 μM) of PF74 compete with host factors for binding to CA (28, 29, 37, 60). Our lead compounds demonstrated effects on both early and late-stage viral replication, similar to PF74 (Table 2) (25) and LEN.

LEN has been observed in both *in vitro* assays and cell-based assays to disrupt core integrity and hyperstabilize the capsid lattice, leading to failure of capsid uncoating and further downstream functions (35, 43, 58, 61, 62). In addition, LEN has been observed to increase levels of viral cores in the cytoplasm and docked at the nuclear pore, and inhibited formation of IN puncta in the nucleus, suggesting that LEN-induced alterations in the capsid lattice block nuclear import (35, 61). In contrast, PF74 has been shown to prevent docking of the viral cores at the nuclear pore, suggesting a different mechanism of blocking nuclear import than LEN (61). PF74 has been shown to stabilize capsids and prevent reverse transcription (63), although some studies have reported PF74 destabilization of the capsid (26). Using the CypA-dsRed assay, we demonstrated that our lead compounds all stabilize native HIV-1 capsid core (Figure 2). Interestingly, our lead compound, ZW-1261, in the fate-of-the-capsid assay appears to destabilize the core at lower concentrations (0.5-1 µM) like PF74; however, at higher concentrations (2.5-5 µM), stabilization of the core was observed (Figure 3B *vs.* 3C), suggesting a possible dual-phase inhibition mechanism at different concentrations. In our nuclear import assay, at low concentrations of ZW-1261 there is an observable decrease in the entry of capsid into the nucleus, while at high concentrations increased entry of capsid into the nucleus is observed (Figure 4A *vs.* 4B). These results correlate with the increase in core stability determined in the fate-of-the-capsid data shown in Figure 3.

It has been reported that HIV-1 infection re-localizes CPSF6 to nuclear speckles that colocalize with the marker SC35 (40, 42). PF74 and GS-CA1 have been reported to prevent the formation of CPSF6 accumulation in nuclear speckles during infection (7, 40, 64). On the other hand, once CPSF6 complexes are already formed in nuclear speckles, PF74 promotes the disassembly of CPSF6 from the nuclear speckles, while GS-CA1 did not affect the preformed CPSF6 complexes (43). Our current results indicate that our lead compound ZW-1261 acts similar to PF74, but different than GS-CA1, to suppress HIV-1 infection (Figure 5).

In the “FG”-binding pocket, several residues are important for host factor and PF74 binding. Residues N57, Q63, and K70 of the CA_NTD_ form hydrogen bonds with PF74, and R173 in the CTD of the neighboring CA monomer forms a cation-π stacking interaction with the indole ring (R3) of PF74 (12, 28, 29). Owing to its large size, LEN participates in several additional interactions that help to stabilize its binding mode in the capsid lattice, including seven hydrogen bonds and 2 cation-π interactions (19, 35). We hypothesize that increased stability of the capsid lattice can be achieved by increasing interactions between CA-targeting molecules like PF74 and LEN at the “FG”-binding site with the CTD of the neighboring CA monomer. Our lead molecule, ZW-1261 is able to engage in increased interactions with the CTD compared to PF74, primarily through its 5-OH modification on R3. This moiety enables interaction of ZW-1261 with K182 and Q179 interaction mediated through an ordered water molecule, which could explain this compound’s increased binding to CA hexamer and antiviral potency compared to PF74. Interestingly, addition of a bulky isopropyl group at the N1-position of R3 in ZW-1527 necessitates a significant change in the R3 binding at the “FG” site. This change in binding positions the isopropyl group closer toward the CTD of the neighboring CA monomer. While the isopropyl group of ZW-1527 itself does not interact with the CTD, this binding mode presents novel opportunities to design compounds with additional modifications at the N1-isoproyl site to form hydrogen bonds or other interactions with the CTD to increase stability of the CA lattice.

Overall, our results reveal that changes to the chemical structure of the R3 ring of PF74 have different effects in binding of analogs to the HIV capsid lattice. This information is useful for the design of future CA-targeting antivirals.

## Materials and Methods

### Cell lines

HeLa P4 MAGI CCR5+ (P4R5) (65) and MT-2 (66, 67) cells were obtained through the AIDS Research and Reference Reagent Program, Division of AIDS, NIAID, NIH. TZM-GFP were generously provided by Dr. Massimo Pizzato, University of Trento (68). HEK 293FT cells were bought commercially (Thermo Fisher Scientific). TZM-GFP is a reporter HeLa cell line that expresses the GFP gene driven by the HIV-1 LTR promoter. GFP expression is activated by Tat expression following HIV infection. All adherent cells were maintained in Dulbecco’s Modified Eagle Medium (DMEM) supplemented with 10% fetal bovine serum (FBS). Suspension cells were maintained in RPMI supplemented with heat inactivated 10% FBS. Cells were incubated at 37°C with 5% CO_2_.

### HIV-1 molecular clones

HIV-1 infectious clones, containing the gag/pol coding regions of primary isolates from subtypes B, C, AE, and AG in an NL4-3 backbone (NL4-3 original clone from the NIH AIDS Research and Reference Reagent Program, (69)) were obtained from Drs. Ujjwal Neogi and Anders Sönnerborg. The HIV-2 subtype B molecular clone plasmid pST (catalog #12444) was obtained from the NIH AIDS Research and Reference Reagent Program (70, 71). Viral stocks of HIV-1 subtypes and HIV-2 were generated by transfection of the molecular clone plasmid into 293T cells, as previously described (72, 73). For expansion of replication-competent viruses, a T75 flask containing 1.8 x 10^6^ MT-2 cells was infected with 2 mL filtered virus, and monitored until syncytia formation was observed, typically 3-5 days. To prepare the expanded viral supernatant for concentration, media was collected, centrifuged for 5 min at 2,000 rcf and filtered through 0.22 µm filter. Viruses were precipitated from the supernatant by incubation with PEG 8,000 (80 g/L) overnight at 4 °C, followed by centrifugation for 40 min at 3,500 rpm. The resulting virus-containing pellet was concentrated 10-fold by resuspension in 1/10^th^ the starting volume of DMEM without FBS and stored at −80 °C.

Vesicular stomatitis virus G protein (VSV-G)-HIV-1_NL4-3_ pseudotyped viruses were generated by co-transfecting envelope-deleted NL4-3 cDNA (pNL4-3ΔEnv) with a VSV-G envelope expression vector. 48 to 72 h post transfection, media was collected, centrifuged for 5 min at 2,000 rcf and filtered through a 0.22 μm polyethersulfone (PES) membrane (Millipore) to remove residual cells.

### HIV-1 capsid targeting molecules

Lenacapavir was purchased from MedChemExpress (catalog # HY-111964). PF74 and all ZW analogs were synthesized by Lei Wang and Zhengqiang Wang at the University of Minnesota as previously described (31, 32). GS-CA1 was generously provided by Stephen Yant (Gilead Sciences, CA).

### Virus Production-Infectivity Assays

To produce virus in the presence of compound, assays were carried out as previously described with modifications (74). HEK 293FT cells were plated at 5 x 10^4^ cells per well in a poly-L-lysine-coated 96-well microtiter plate. Twenty-four hours later, cells were transfected with 0.1 µg of full length pNL4-3 plasmid per well using FuGENE® HD Transfection Reagent (Promega). Dilutions of compounds and controls were then added to transfected cells 6 h after transfection. The supernatants were harvested 72 h after transfection. One microliter of the transfected supernatants was used for subsequent infection (100-fold dilution) in a fresh 96-well microtiter plate of TZM-GFP cells plated at 1 x10^4^ cells per well. HIV-1 replication was assessed 48 hours post infection (hpi) by counting the number of GFP positive cells on a Cytation-5 Imaging Reader (BioTek) with GFP filter cubes and 4x objective image montage. Compound-induced effects manifested as a decrease in infectivity in the target cells (measured as GFP cell count), normalized against the infectivity of virus produced from DMSO-treated cells. For dose responses, values were plotted in GraphPad Prism and analyzed with the log (inhibitor) *vs.* normalized response – variable slope equation to obtain the EC_50_ value.

### Early vs. late-stage experiments

To evaluate the antiviral activity of compounds, TZM-GFP cells were plated at 1×10^4^ cells per well in a 96-well microtiter plate. Twenty-four hours later, cells were infected with WT HIV-1_NL4-3_ (MOI = 1) in the presence of compound for 48 h. Compound-treated wells and DMSO vehicle controls contained 1% final concentration of DMSO and 1 µg/mL final concentration of DEAE-dextran. HIV-1 infection was assessed by counting the number of GFP positive cells on a Cytation-5 Imaging Reader (BioTek) with GFP filter cubes and 4x objective image montage. Compound-induced effects manifested as a decrease in infectivity (measured as count of GFP positive cells), normalized against the infectivity in the DMSO treated (vehicle control) cells. For dose responses, values were plotted in GraphPad Prism and analyzed with the log (inhibitor) *vs.* normalized response – variable slope equation to obtain the half maximal effective concentration (EC_50_). Single-cycle infections assays were performed and graphed as described above except with VSV-G-HIV-1_NL4-3_ pseudotyped virus.

### HIV-1 capsid stability assays

HIV-1 capsid stability measurements were done as described previously (33, 34). Briefly, pseudoviruses co-labeled with integrase-superfolder-GFP (INsfGFP) marker of internal vRNPs and CypA-DsRed (CDR) marker of the outer capsid lattice, were adhered to poly-l-lysine treated 8-well chambered slides for 30 minutes at 4°C. To measure the loss of CDR, virions were permeabilized with Saponin (100 μg/ml) in the presence of DMSO or 10 µM PF74, ZW-1559, ZW-1261, ZW-1514, ZW-1517, or ZW-1524. Single-plane time-lapse images from 4 independent fields of view were acquired every 30 seconds for 60 minutes using a Zeiss LSM780 or Leica SP8 confocal microscope. Laser lines 488, and 561 nm lasers was used to visualize INsfGFP (green), and CDR (red) puncta, respectively. Images were analyzed offline using the spot detection algorithm of the ICY image analysis software, and the average number of INsfGFP and CDR spots per field of view over time was determined and plotted.

### Fate-of-the-capsid assay

The fate-of-the-capsid assay was performed as previously described (36, 38, 40). Human A549 cells were infected with p24-normalized amounts of WT HIV-1-GFP virus in the presence of DMSO as vehicle control, 10 µM PF74 control, and lead compound ZW-1261 at concentrations of 0.5, 1, 2.5, and 5 µM. After incubation at 37 °C for 12 hours, cells were detached with 7 mg/mL Pronase for 5 minutes on ice and washed 3 times with ice-cold PBS. Cell pellets were resuspended in hypotonic buffer (10 mM Tris-HCl, pH 8.0; 10 mM KCl; 1 mM EDTA) and incubated for 20 minutes on ice. Cells were lysed in a 7.0 mL Dounce homogenize. Cellular debris and nuclear fraction were removed by centrifugation for 7 minutes at 3,000 rpm. The supernatant fraction was layered onto a 50% sucrose (weight:volume) cushion in 1x PBS and centrifuged at 125,000 x g for 2 hours at 4 °C in a Beckman SW41 rotor. Input, soluble, and pellet fractions were analyzed by western blotting using anti-p24 antibody.

### Nuclear import assay

The nuclear import assay was performed as previously described (40, 41). Cells (5 × 10^6^ cells) were challenged with HIV-1 viruses at a MOI = 2 for different time points. Cells were harvested using trypsin for 2–5 min at 37 °C. Harvested cells were washed twice with 1x cold PBS by centrifugation at 2000 rpm for 7 min at 37 °C. Supernatant was discarded and cell pellet was resuspended in 1 mL of PBS. 1/10 aliquot of the cell suspension (100 μL) was centrifuged (2000 rpm for 7 minutes at 37 °C), supernatant was discarded and cell pellet was resuspended in 35 μL of WCE (50 mM Tris pH = 8.0, 280 mM NaCl, 10% glycerol, 0.5% NP-40, 5 mM MgCl_2_, 250 units/mL Benzonase, 1x protease inhibitor), incubated for 1 hr on ice, centrifuged at 13400 rpm for 1 hour at 37 °C, then resulting supernatant was mixed with 5x Laemmli buffer and used to measure the total amount of capsid. The rest of the cell suspension (900 μL) was centrifuged (2000 rpm for 7 min at 37 °C), supernatant was discarded and cell pellet was resuspended in 315 μL of lysis buffer (10 mM Tris pH = 6.8, 1 mM DTT, 1 mM MgCl_2_, 10% sucrose, 100 mM NaCl, 0.5% NP-40, 1x protease inhibitor) and incubated for 5 min on ice. The sample was then centrifuged at 2500 rpm for 2 min at 37 °C. The resulting supernatant and pellet correspond to cytosolic and nuclear fractions, respectively. Next, 1/9 aliquot of supernatant (35 μL) was mixed with 5x Laemmli buffer and used as cytosolic fraction. The nuclear pellet was washed twice using 1 mL of lysis buffer without NP-40 (10 mM Tris, pH = 6.8, 1 mM DTT, 1 mM MgCl_2_, 10% sucrose, 100 mM NaCl, 1x protease inhibitor) by gently inverting the tube 2–3 times. The sample was then centrifuged at 2500 rpm for 2 min at 37 °C. The nuclear pellet was resuspended in 315 μL of extraction buffer (10 mM Tris, pH = 6.8, 1 mM DTT, 1 mM MgCl_2_, 10% sucrose, 400 mM NaCl, 1x protease inhibitor), and incubated on ice for 10 min. Subsequently, the sample was centrifuged at 7000 rpm for 2 min at 37 °C. 1/9 aliquot of the supernatant (35 μL) was mixed with 5x Laemmli buffer and used as nuclear fraction. Proportional amounts of total, cytosolic, and nuclear fractions were analyzed by western blot using anti-p24, anti-Nopp140, anti-α-tubulin, or anti-GAPDH antibodies.

### Immunofluorescence microscopy

Immunofluorescence microscopy assays were performed as previously described (43). Human HeLa cells were seeded on poly-L-lysine–coated chamber slides (BD Biosciences) and regular chamber slides (Nalgene Nunc) and infected with HIV-1 virus-like particles in presence of indicated compounds. Chamber slides were then washed with PBS and incubated with paraformaldehyde (4% PFA, 1x PBS). Cells were incubated first with 0.1% Triton X-100 for 5 min for cell permeabilization and then with blocking solution (3% BSA, 13 PBS) for 30 min to prevent non-specific binding. Samples were incubated with corresponding antibodies in blocking solution (anti-human CPSF6; anti-SC-35). Cells were washed with 1x PBS and then incubated with the corresponding secondary antibodies in blocking solution. Chamber slides were mounted using Gel Mount (Biomedia) and imaged using a Deltavision epifluorescent microscope system fitted with an automated stage (Applied Precision, Inc). Images were captured in z series on a CCD digital camera using a 63X lens. Out-of-focus images were digitally removed using the Softworks deconvolution software (Applied Precision, Inc).

### Biolayer Interferometry (BLI)

BLI experiments were performed as previously described (44, 45). Briefly, Octet® biosensors containing Qiagen Penta-HIS antibodies (HIS1K biosensors, Sartorius) were allowed to equilibrate in running buffer, 20 mM Tris pH 8.0, 40 mM NaCl, 20 mM Imidazole, 0.6 M Sucrose, and BSA (1%), for 15 minutes. C-terminal His-tagged CA121 crosslinked hexamers expressed and purified as previously described (44, 45) were allowed to bind to the biosensors for 600 s at 100 µg/mL. After two baseline steps, the protein-loaded biosensors were introduced to the compounds at 20, 10, and 5 µM for 50 s. Dissociation was performed in a new well for 100 s. For LEN experiments, the protein was loaded at 200 µg/mL for 300 s. Additionally, the association and dissociation time were changed to 200 and 2700 s, respectively. The experiments used double reference subtraction and Savitzky-Golay Filtering. The Y-axis was aligned to either the baseline or dissociation step, and the inter-step correction was set to the baseline. Analysis was performed using the Octet® Analysis Studio software.

### Crystallization, data collection, and structure determination

Native, full-length (FL) WT HIV-1 CA in a pET11a vector was expressed and purified, as previously described (12). Hexagonal CA crystals grew in specific conditions, as previously described (12, 34). CA crystals were soaked in a solution containing ZW-1261, ZW-1514, or ZW-1527 (1.25 mM final concentration with 5% DMSO) for ∼4 h, then briefly transferred to a solution containing 22% glycerol for cryoprotection before flash freezing in liquid nitrogen.

Data were collected on a Dectris Eiger 16 M detector at Advanced Photon Source (APS) beamline 22-ID at the Argonne National Laboratory. For the WT HIV CA FL/ZW-1261 complex, data were processed to 2.7 Å using XDS (74), and indexed in hexagonal space group P6 with unit cell dimensions *a*, *b* = 90.9 Å and *c* = 56.1 Å, and one CA molecule in the asymmetric unit. For the WT HIV CA FL/ZW-1514 complex, data were processed to 2.8 Å using XDS (74), and indexed in hexagonal space group P6 with unit cell dimensions *a*, *b* = 92.3 Å and *c* = 56.9 Å, and one CA molecule in the asymmetric unit. For the WT HIV CA FL/ZW-1527 complex, data were processed to 2.5 Å using XDS (74), and indexed in hexagonal space group P6 with unit cell dimensions *a*, *b* = 92.7 Å and *c* = 57.4 Å, and one CA molecule in the asymmetric unit. Each dataset was analyzed using XTRIAGE, which determined that no twinning was present in any of the datasets (75–77). Initial phases were solved via molecular replacement with Phaser (78), using coordinates of a native full-length CA in complex with PF74 (PDB ID: 4XFZ) (12) as a starting model, with all ligands and solvent removed. The resulting model was refined using REFMAC5 (79). The coordinates and ligand topology of ZW-1261, ZW-1514, and ZW-1527 were generated using PRODRG in the CCP4 Suite (80, 81) or the Phenix electronic Ligand Builder and Optimization Workbench (eLBOW) (76, 82). ZW-1261, ZW-1514, and ZW-1527 were built into their respective models using difference Fourier maps calculated in the absence of ligand. The CA/ZW-1261, CA/ZW-1514, and CA/ZW-1527 models were improved through several iterative rounds of model building and refinement using Coot (83) and REFMAC5 (79), respectively. The final models were validated using MOLPROBITY (84, 85). The final structure factors and coordinates have been deposited into the Protein Data Bank and are available under accession codes 7M9F (WT HIV CA FL in complex with ZW-1261), 9Z9Z (WT CA FL in complex with ZW-1514), and 9ZA0 (WT CA FL in complex with ZW-1527). Data collection, processing, and refinement statistics are provided in Supplementary Table S1.

## Acknowledgements

The following reagents were obtained through the NIH HIV Reagent Program, Division of AIDS, NIAID, NIH: Human Immunodeficiency Virus Type 2 (HIV-2) ST Infectious Molecular Clone, ARP-12444, contributed by Dr. Beatrice Hahn and Dr. George Shaw; P4 MAGI CCR5+ Cells, ARP-3580, contributed by Dr. Nathaniel Landau, Aaron Diamond AIDS Research Center, The Rockefeller University; and Human T-Lymphotropic Virus Type 1 (HTLV-1)-Infected MT-2 Cells, ARP-237, contributed by Dr. Douglas Richman.

This research was supported in part by the National Institutes of Health (R01 AI120860 and P30 AI050409 to S.G.S.; R21 AI189247 to K.A.K.; F31 AI174951 to W.M.M.; F31 AI172618 to A.A.S.; F31 AI94923 to S.M.R.; W.M.M. and A.A.S. were supported in part by T32 GM135060). The content is solely the responsibility of the authors and does not necessarily represent the official views of the National Institutes of Health. S.G.S. acknowledges the Nahmias-Schinazi Distinguished Chair in Research.

X-ray data were collected at the Southeast Regional Collaborative Access Team (SER-CAT) 22-ID beamline at the Advanced Photon Source, Argonne National Laboratory. SER-CAT is supported by its member institutions, and equipment grants (S10 RR25528, S10 RR028976 and S10 OD027000) from the National Institutes of Health. The Advanced Photon Source (APS), a U.S. Department of Energy (DOE) Office of Science User Facility is operated for the DOE Office of Science by Argonne National Laboratory under Contract No. DE-AC02-06CH11357.

## Author contributions

Conceptualization, K.A.K, Z.W. and S.G.S.; methodology & investigation, K.A.K, W.M.M., L.W., H.D., H.Z., A.E.C., Z.C.L., A.H., A.N., A.C.F., C.L., M.E.C., A.A.S., S.M.R., S.C. and P.R.T.; visualization, K.A.K., W.M.M., A.C.F. and F.D.-G.; resources, Z.W., A.C.F., G.B.M., F.D.-G. and S.G.S.; writing – original draft, K.A.K., and W.M.M.; writing – review & editing W.M.M., K.A.K., P.R.T., A.C.F., F.D.-G. and S.G.S.; funding acquisition, K.A.K., Z.W., and S.G.S..

