## Supplemental Table 1 for "Structural, biophysical, and virological mechanistic characterization of HIV-1 capsid-targeting antivirals"

**Table S1. X-ray data collection and refinement statistics**

| | WT CA FL / ZW-1261 PDB: 7M9F[a] | WT CA FL / ZW-1514 PDB: 9Z9Z | WT CA FL / ZW-1527 PDB: 9ZA0 |
|---|---|---|---|
| **Data collection** | | | |
| Beamline | APS 22-ID | APS 22-ID | APS 22-ID |
| Wavelength (Å) | 1.00000 | 1.00000 | 1.00000 |
| Resolution (Å) | 2.70 (2.70-2.77)[b] | 2.81 (2.81-2.98)[b] | 2.49 (2.49-2.64)[b] |
| Space group | *P*6 | *P*6 | *P*6 |
| Cell dimensions | | | |
| *a*, *c* (Å) | 90.9, 56.1 | 92.3, 56.9 | 92.7, 57.4 |
| Observed reflections | 52,178 | 68,541 | 102,959 |
| Unique reflections | 7,239 | 13,215 | 9,941 |
| Redundancy | 7.2 (6.1) | 5.2 (4.8) | 10.3 (10.7) |
| Completeness (%) | 97.9 (91.1) | 99.6 (98.6) | 100 (100) |
| $R_{meas}$[c] | 0.118 (0.84) | 0.079 (0.76) | 0.071 (1.22) |
| $CC_{1/2}$ | 99.6 (78.3) | 99.8 (77.6) | 99.8 (73.7) |
| Avg I/σ | 13.6 (3.4) | 12.5 (1.5) | 17.5 (2.1) |
| **Refinement statistics** | | | |
| Resolution (Å) | 45.71-2.70 | 79.92-2.81 | 80.28-2.49 |
| No. of reflections (working) | 6,881 | 6,503 | 9,465 |
| No. of reflections (test) | 356 | 360 | 472 |
| $R_{work}$[d] | 0.192 | 0.248 | 0.268 |
| $R_{free}$[e] | 0.250 | 0.319 | 0.300 |
| Overall B value (Å$^2$) | 71.0 | 111.5 | 91.2 |
| Wilson B value (Å$^2$) | 63.5 | 96.5 | 75.4 |
| Ramachandran plot (%)[f] | | | |
| Favored | 97.3 | 90 | 97 |
| Allowed | 2.7 | 7 | 3 |
| Disallowed | 0 | 2 | 0 |
| All-atom clashscore | 4 | 10 | 5 |
| RMSD Bond length (Å) | 0.018 | 0.005 | 0.007 |
| RMSD Angle (°) | 1.978 | 1.266 | 1.299 |

[a] Data collection and refinement statistics for 7M9F reported in Sun, Q., *et al.* 2021, *Viruses* 13:920.

[b] Values in parentheses are for the outer resolution shell.

[c] $R_{meas} = \Sigma_{hkl} \sqrt{\frac{n}{n-1}} \sum_{j=1}^{n} |I_{hkl,j} - \langle I_{hkl} \rangle| \;/\; \Sigma_{hkl} \Sigma_j I_{hkl,j}$.

[d] $R_{work} = \Sigma_{hkl} |F_{obs} - F_{calc}| \;/\; \Sigma_{hkl} |F_{obs}|$.

[e] $R_{free} = R_{work}$, except 5% of the data excluded from the refinement.

[f] Evaluated by MolProbity (Davis, I. W., *et al.* 2007. *Nucleic Acids Res* 35:W375-383).
